# Out-of-equilibrium gene expression fluctuations in presence of extrinsic noise

**DOI:** 10.1101/2023.02.14.528039

**Authors:** Marta Biondo, Abhyudai Singh, Michele Caselle, Matteo Osella

**Affiliations:** Department of Physics, University of Turin and INFN, via P. Giuria 1, I-10125 Turin, Italy; Department of Electrical and Computer Engineering, University of Delaware, Newark, DE 19716, USA

## Abstract

Cell-to-cell variability in protein concentrations is strongly affected by extrinsic noise, especially for highly expressed genes. Extrinsic noise can be due to fluctuations of several possible cellular factors connected to cell physiology and to the level of key enzymes in the expression process. However, how to identify the predominant sources of extrinsic noise in a biological system is still an open question. This work considers a general stochastic model of gene expression with extrinsic noise represented as colored fluctuations of the different model rates, and focuses on the out-of-equilibrium expression dynamics. Combining analytical calculations with stochastic simulations, we fully characterize how extrinsic noise shapes the protein variability during gene activation or inactivation, depending on the prevailing source of extrinsic variability, on its intensity and timescale. In particular, we show that qualitatively different noise profiles can be identified depending on which are the fluctuating parameters. This indicates an experimentally accessible way to pinpoint the dominant sources of extrinsic noise using time-coarse experiments.

**Author summary:** Genetically identical cells living in the same environment may differ in their phenotypic traits. These differences originate from the inherent stochasticity in all cellular processes, starting from the basic process of gene expression. At this level, large part of the variability comes from cell-to-cell differences in the rates of the molecular reactions due to stochasticity in the level of key enzymes or in physiological parameters such as cell volume or growth rate. Which expression rates are predominantly affected by these so-called “extrinsic” fluctuations and how they impact the level of protein concentration are still open research questions. In this work, we tackle the protein fluctuation dynamics while approaching a steady state after gene activation or repression in presence of extrinsic noise. Our analytical results and simulations show the different consequences of alternative dominant sources of extrinsic noise, thus providing an experimentally-accessible way to distinguish them in specific systems.

## Introduction

Cellular processes are subjected to stochastic fluctuations. These fluctuations (or noise) can lead to phenotypic differences even in genetically identical cells sharing the same history and environment. Noise can often be detrimental for the cell since it affects the precision and reliability of several processes, for example related to signalling. Indeed, a high noise level has been associated to partial or complete loss of cellular functions [1–3], and there is evidence of evolutionary selection against cellular noise [4, 5]. On the other hand, molecular noise can be beneficial in other circumstances [6]. It can be exploited to drive genetically identical cells to different cell fates in multi-cellular organisms [7–13], or it can induce the phenotypic diversification at the basis of bet-hedging strategies that protect microbial cell populations from sudden environmental changes [14–17].

Focusing specifically on the gene expression process, two possible sources of fluctuations can be defined, i.e., intrinsic and extrinsic noise. Intrinsic noise arises from the inherently stochastic nature of the molecular reactions involved in transcription, translation, and degradation of messenger RNAs (mRNAs) and proteins. Extrinsic noise is instead the result of fluctuations in global cellular factors such as the concentration of key enzymes (e.g., ribosomes and polymerases) involved in the process [18, 19]. These extrinsic fluctuations can also arise from cell-to-cell differences in metabolic states [20], cellular signalling [21, 22], cell-cycle stage [23–25], and general cell physiology (i.e., cell growth rate, doubling time, volume, etc.) [26–32]. Noise propagation through gene regulation is another key source of gene expression noise. In fact, the number of regulatory inputs of a gene is correlated with its expression variability in *E. coli* [33] and eukaryotes [34, 35], and the topology of the regulatory network can play a crucial role in the extent of noise propagation [36, 37].

A large amount of theoretical and experimental work has focused on disentangling the contributions to protein fluctuations from intrinsic and extrinsic sources (see for example ref. [38–40]). On the other hand, what are precisely the dominant sources of extrinsic noise, and thus how they have to be correctly inserted in effective models of gene expression are still open questions, even if extrinsic noise actually seems the main noise source for sufficiently highly expressed genes [35, 41, 42].

From a modelling standpoint, extrinsic noise can be defined as fluctuations of the parameters of the gene expression process such as degradation and production rates. All the possible biological sources of extrinsic noise listed above can affect in complex ways one or more parameters. For example, growth rate fluctuations directly impact the dilution rate of proteins through volume fluctuations, but at the same time can change the protein production rates by affecting the concentration of key enzymes, such as ribosomes, that are closely coupled with cell growth [43]. The problem is to pinpoint which are the parameters that are mostly affected in the biological system of interest, in order to design the correct minimal model of the stochastic process of gene expression.

This work focuses on this problem by looking at the consequences on the out-of-equilibrium dynamics of gene expression when extrinsic noise affects different rates of production or degradation. Specifically, we will provide analytical expressions supported by simulations that fully characterize how the protein level and its fluctuations evolve during gene activation and inactivation in presence of extrinsic fluctuations on different parameters. On top of the theoretical interest of this analysis, the results have immediate practical applications. In fact, the different dynamic profiles of protein noise that we characterized naturally provide a relatively simple method to experimentally distinguish between different extrinsic-noise scenarios.

## Materials and methods

### An effective model of stochastic gene expression with extrinsic noise

The “standard model” of stochastic gene expression takes into account messenger RNA and protein production and degradation as first-order chemical reactions [44–47]. After activation, the gene is transcribed by RNA polymerases in mRNAs with a fixed rate *k*_*m*_; each mRNA (*m*) is in turn translated into proteins (*p*) with rate *k*_*p*_. Proteins and mRNAs are also removed at specific constant rates (*γ*_*m*_ and *γ*_*p*_). Fig. 1 schematically represents this set of reactions. The degradation rate of mRNA molecules sets their average lifetime (1/*γ*_*m*_), which is typically short compared to the average residence time of proteins (1/*γ*_*p*_). Especially in microorganisms, the average lifetime of mRNAs is just few minutes [41]. On the other hand, the rate *γ*_*p*_ is mainly set by dilution due to cell growth and division [41, 47–49], and thus the protein lifetime is essentially set by the cell doubling time *τ*_*p*_ = ln(2)/*γ*_*p*_ [50].

**Fig 1.**
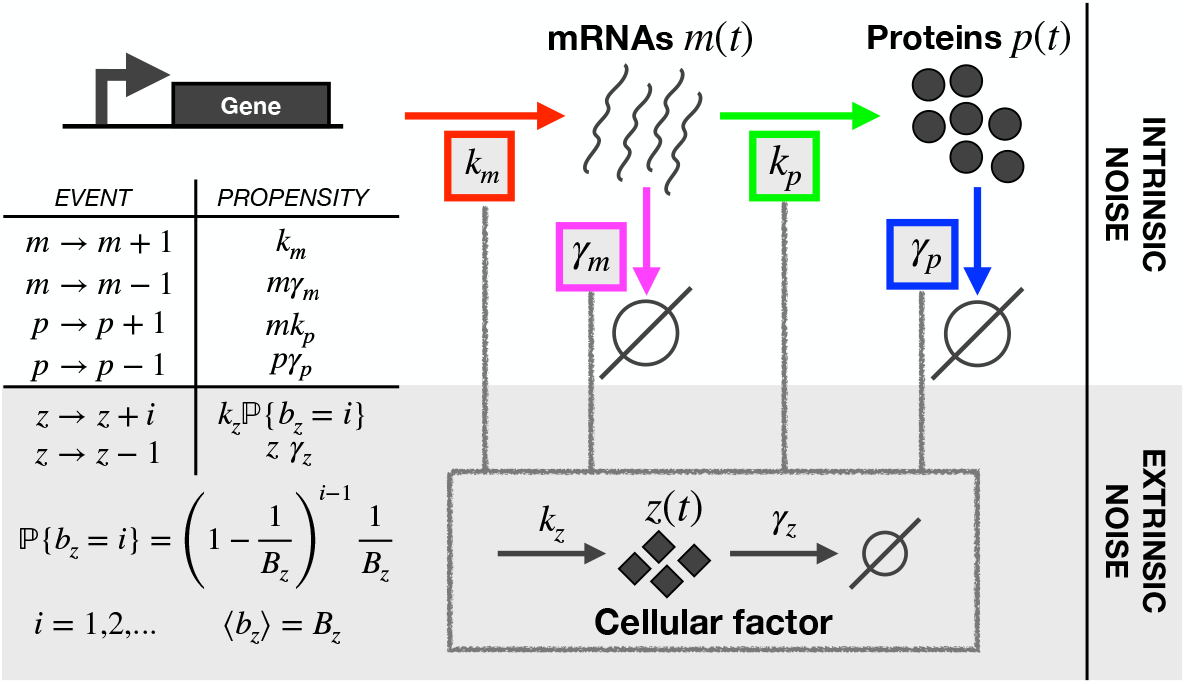
Model of the gene expression process with extrinsic noise. Extrinsic noise is included in a basic two-step model of stochastic gene expression by introducing the cellular factor *z*(*t*). *z*(*t*) can affect any parameter of the model: transcription and translation rates (*k*_*m*_ and *k*_*p*_), as well as degradation rates for mRNAs and proteins (*γ*_*m*_ and *γ*_*p*_). The table on the left lists the possible reactions for mRNAs *m*, proteins *p* and for the cellular factor *z* with the respective propensity functions that set the event frequencies.

The master equation describing this simple two-step model of gene expression (Fig. 1) can be solved assuming that the promoter is activated at time *t* = 0 and the initial number of mRNAs and proteins is zero [47]. In particular, the dynamics of the average protein numbers is described by

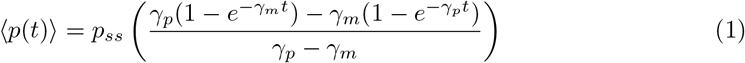

where *p*_*ss*_ = *k*_*m*_*k*_*p*_/*γ*_*m*_*γ*_*p*_ is the average protein level at steady state. The noise can be quantified by the Coefficient of Variation 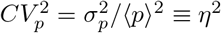. For stable proteins, the timescale separation between the dynamics of mRNAs and proteins (*γ*_*m*_ ≫ *γ*_*p*_) can be used to derive a compact expression for the time evolution of the intrinsic noise [47] and for its equilibrium value:

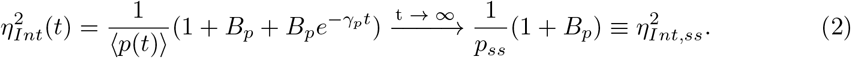

*B*_*p*_ = *k*_*p*_/*γ*_*m*_ is the protein burst size, i.e., the average number of proteins produced by a single mRNA during its lifetime. As the expression at steady state shows, burstiness introduces an amplification factor with respect to Poisson noise. We focus on realistic and relatively high levels of expression (*p*_*ss*_=2000 proteins in most examples we will describe) and sufficiently low protein burst sizes (few units), to focus on the quite common situation in which the intrinsic contribution to expression noise is not dominant with respect to extrinsic part, at least at steady state.

We will also consider the dynamics of a gene at steady state that is inactivated at the transcriptional level. In this case, the average protein evolution in time is given by

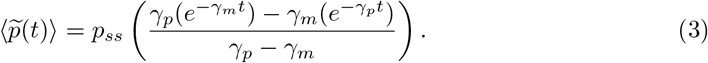

So far we have only included intrinsic fluctuations since all the rates were constant. To include extrinsic fluctuations, we introduce a generic cellular factor *z* that can affect production or degradation rates (Fig. 1). For example, fluctuations in RNA polymerases or ribosomes will be captured by a direct action of the factor *z* on the production rates *k*_*m*_ or *k*_*p*_. More generally, the cellular factor *z* can capture the consequences that fluctuations in cell physiology can have on gene expression by modulating the affected rates.

Extrinsic noise is typically a colored noise [18, 41] with a timescale that depends on the effective and typically unknown source of fluctuations. To explore the role of both extrinsic fluctuations strength and timescale, we model the dynamics of the cellular factor as a bursty birth-and-death process with constant production and degradation rates (*k*_*z*_ and *γ*_*z*_). With this modelling choice, production events are geometrically distributed with bursts of average size *B*_*z*_ [39]. Tuning *k*_*z*_*, γ*_*z*_ and *B*_*z*_, the extent (*CV*_*z*_) and the timescale (*τ*_*z*_ = ln(2)/*γ*_*z*_) of extrinsic fluctuations can be independently modulated.

We can thus introduce extrinsic noise on any biochemical rate by multiplying its value to the cellular factor. In other words, we can substitute any parameter *θ* of the system with 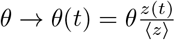. In this way, the average parameter value is still *θ*, i.e., the value set in absence of extrinsic fluctuations, but it fluctuates according to *z*.

Intrinsic noise *η*_*Int*_ will be defined as the variability associated with the model of stochastic gene expression with constant parameters. Instead, extrinsic noise *η*_*Ext*_ can be quantified as the difference between the total measured protein variability and the intrinsic part.

### Assessment of the time evolution of protein cell-to-cell variability

The definition of intrinsic and extrinsic noise naturally implies the decomposition 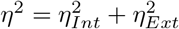 [51]. This decomposition is also valid out-of-equilibrium and for each parameter *θ* affected by extrinsic fluctuations of strength defined by *CV*_*z*_ and timescale by *τ*_*z*_. Therefore, we can write

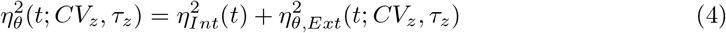

We are interested in the dynamics of gene-expression noise approaching a steady state. In the case of fluctuations of the production rates *k*_*m*_ or *k*_*p*_, it is possible to derive the exact transient protein noise expression by solving the corresponding system of ordinary differential equations. The details of the calculation can be found in the SI File, but the main result is that for extrinsic noise acting on production rates we can provide an analytical estimate of the time evolution of protein noise.

Unfortunately, the same approach cannot be applied when extrinsic fluctuations affect the dilution rate *γ*_*p*_. Nevertheless, an approximate expression for the steady-state protein noise 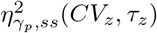 can still be calculated as a function of the timescale and strength of extrinsic noise. In order to provide also an expression for the out-of-equilibrium noise, we can assume that the time dependence in the noise expression can be factorized as

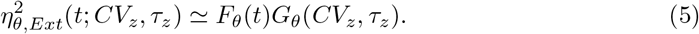

*F*_*θ*_(*t*) explicitly captures the time-dependent part of the impact of *θ* fluctuations on gene expression noise, while *G*_*θ*_(*CV*_*z*_*, τ*_*z*_) is time independent and takes into account the features of extrinsic noise. Assuming this factorization, an intuitive although approximate expression can be provided using the framework of sensitivity analysis. Basically, we can first evaluate the fluctuations due to the extrinsic factor acting on the parameter *θ* at steady state, and this is possible for every choice of *θ*. We can then further assume that the effect of the fluctuating parameter *θ* on the protein level is predominantly set by the first-order dependency of the average protein dynamics on *θ*. In other words, we can estimate the protein variance in time using the propagation of uncertainty as

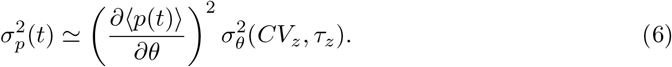

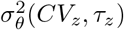 is the variance of the parameter *θ*, which is simply given by the variance of the extrinsic factor *z*. The relation can be rephrased for the Coefficient of Variation as

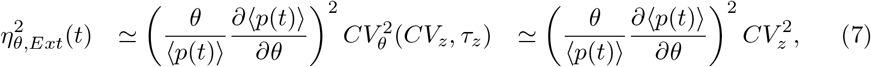

where, by definition, 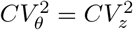 at any given time.

This estimate provides the functional dependency of *F*_*θ*_(*t*), while the time-independent noise 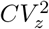 can be included in the factor *G*_*θ*_(*CV*_*z*_*, τ*_*z*_) of the decomposition in Eq. 5. The approximation considers *p*(*t*) and *θ* as continuous variables and neglects the impact of the strength and timescale of extrinsic fluctuations on the time dependent factor.

From the expression of *p*(*t*), it is easy to show that lim_*t*→∞_ *F*_*θ*_(*t*)=1 for any parameter *θ*. This crucial observation implies that, in order to have consistency at steady state, the factor *G*_*θ*_(*CV*_*z*_*, τ*_*z*_) has to be equal to the extrinsic noise at steady state 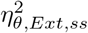.

Therefore, we finally have explicitly defined the two factors of Eq 5 as

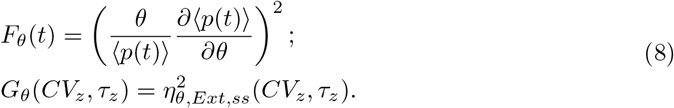

As discussed above, analytical expressions can be calculated for the total protein noise at steady state 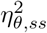 as a function of the strength and the timescale of extrinsic fluctuations (see Eqs. S15, S18, S22). Analogously, the intrinsic noise 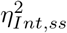 can be calculated [47]. Therefore, the extrinsic part can be extracted with a simple subtraction from 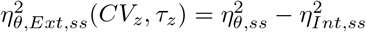.

All the analytical expressions have been tested with extensive numerical simulations using the exact Gillespie algorithm [52] as detailed in the SI File (section S2). In the following sections the analytical curves will always be supported by and compared to numerical results.

## Results

### Extrinsic fluctuations of the protein dilution rate alter the average protein dynamics

The first result of this work is to provide analytical expressions for the protein dynamics and its variability in presence of extrinsic fluctuations acting on different gene expression parameters. These analytical results are detailed in the Methods section and in the SI File and will be discussed in the following sections.

Here, we focus on the expected value of the protein level at steady state for different extrinsic noise sources. The different expressions we obtained show that extrinsic fluctuations acting on production rates (i.e., translation rate *k*_*p*_ and transcription rate *k*_*m*_) do not alter the average protein level *p*_*ss*_ described by Eq. 1 at equilibrium, but fluctuations of the dilution rate *γ*_*p*_ can significantly change it (Eq. S21). When the extrinsic noise acts on the dilution rate, the system of differential equations describing the protein moment dynamics is not closed, but an approximate moment-closure technique (see SI File, section 1.3, Eqs. S20) can be used to estimate the expected protein level at steady state as

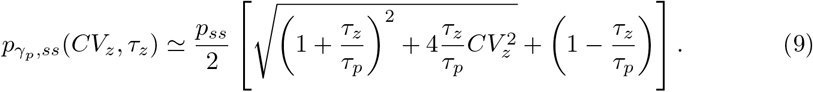

The expression above indicates how the protein level depends on the cellular factor *z* and how it is different by the value *p*_*ss*_ = *k*_*m*_*k*_*p*_/*γ*_*m*_*γ*_*p*_ without extrinsic noise. In particular, it increases with the fluctuation strength *CV*_*z*_ and has a sigmoidal dependence on the fluctuation timescale *τ*_*z*_. These analytical predictions are well supported by stochastic simulations (Supplementary Fig.S2).

Therefore, the noise-induced alteration of the average behaviour, also known as deviant effect [37, 53], generates a discrepancy with respect to the classic deterministic prediction *p*_*ss*_, and this discrepancy grows with the level of extrinsic fluctuations. Thus, deviations from the expected average protein value could in principle be used in experimental settings as hallmarks of large extrinsic fluctuations acting on degradation rates. However, this observation would practically require the knowledge of the process parameter values, which are often not known.

This work aims to fully characterize the complex interplay between extrinsic fluctuations and protein dynamics in single cells. We will show that this characterization is instrumental to define easily measurable signatures of the possible dominant sources of extrinsic noise in a system, even when the specific parameter values are not known. However, in order to do so, we need to compare the expected single-cell protein dynamics with extrinsic fluctuations acting on different parameters with a fixed common average steady-state value. As explained in detail in the SI File (section S1.4) and shown in Fig.S3, we implemented this classic “mathematically controlled comparison” [49, 54] by taking into account the deviant effects and constraining the average steady-state protein value for any magnitude and timescale of extrinsic fluctuations, independently of which the noisy parameter is.

### The dynamics of cell-to-cell variability is strongly dependent on the dominant source of extrinsic noise

This section describes the single-cell expression dynamics when a gene is activated in presence of extrinsic fluctuations. The main observation is that the protein noise dynamics is qualitatively different depending on which parameter is predominantly affected by extrinsic fluctuations. In fact, even if the average protein dynamics is constrained to be the same (Fig. 2A), as explained in the previous section, the protein probability densities are significantly different at short times depending on the extrinsic noise source (Fig. 2B). The out-of-equilibrium protein fluctuations are higher for noise affecting production rates rather than degradation rates. This distinction cannot be done at equilibrium since the probability densities progressively collapse as the protein level approaches the steady state.

**Fig 2.**
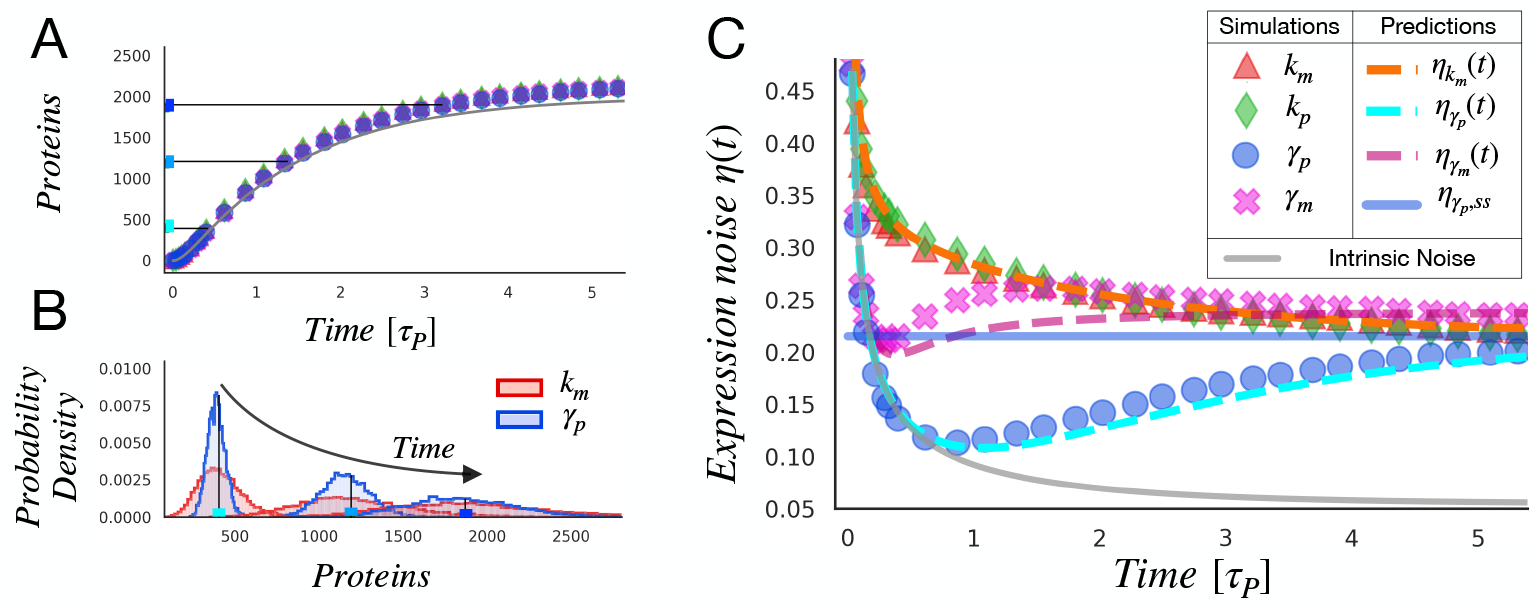
Cell-to-cell variability for different sources of extrinsic noise. We compare the time evolution of expression variability during the activation dynamics when extrinsic fluctuations affect a single rate of production (*k*_*m*_*, k*_*p*_) or of degradation (*γ*_*m*_ or *γ*_*p*_). Even if the extrinsic noise properties are fixed (*CV*_*z*_=0.3, *τ*_*z*_/*τ*_*p*_=1, ⟨*z*⟩ =1000 copies), different protein noise profiles *η*(*t*) can be observed depending on the fluctuating parameter. A) Thanks to the controlled comparison, the average protein level approaches a fixed steady state following Eq. (1), independently of the source of extrinsic noise. B) The protein probability densities are qualitatively different in presence of fluctuations of *k*_*m*_ or *γ*_*p*_. The mean protein levels at different times are reported as vertical lines and correspond to the horizontal lines in (A). C) The total gene expression noise, quantified by the coefficient of variation, is reported as a function of time during the transient regime (time ∈ [0; 5]*τ*_*p*_). The continuous grey line represents the variability without extrinsic noise *CV*_*z*_=0, i.e., only the intrinsic noise described by Eq. (2). The horizontal blue line marks the approximate prediction for the steady-state expression variability under fluctuations of *γ*_*p*_, to which the values of the simulations asymptotically tend. The dashed lines correspond to the theoretical predictions (Eqs. (10) and (14)), which are well compatible with the simulation results (symbols).

Fig. 2C explicitly reports the dynamics of protein expression noise *η*(*t*), measured with the coefficient of variation, and shows its different behaviors depending on the dominant source of extrinsic noise. The trends in presence of fluctuations of the production rates (i.e., *k*_*m*_ or *k*_*p*_) are qualitatively and quantitatively very similar. The simulations are well explained by the exact formula (fully derived in the SI File sections S1.1 S1.2) for the time-evolution of protein noise in the presence of transcription rate fluctuations (orange dashed line in Fig. 2):

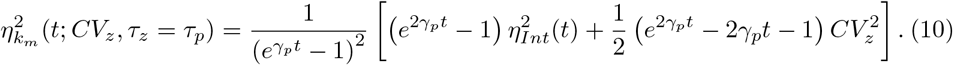

The corresponding steady state limit is given by

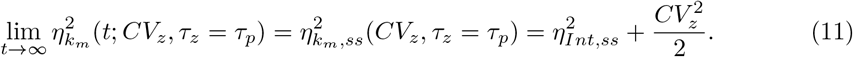

The time dependence of Eq. (10) is analogous to the one of intrinsic fluctuations, given by Eq. (2) and corresponding to the grey line in Fig. 2C. The presence of extrinsic noise essentially increases the relaxation point to the higher noise value predicted by Eq. (11) without changing the monotonous decreasing trend.

This result can be intuitively understood by considering the factorization proposed in Eq. (5). Since *∂*⟨*p*(*t*) ⟩/*∂k*_*m*_ = ⟨*p*(*t*) ⟩/*k*_*m*_ and *∂*⟨*p*(*t*) ⟩/*∂k*_*p*_ = ⟨*p*(*t*) ⟩/*k*_*p*_, the factors 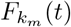 and 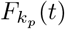 do not depend on time, thus explaining why 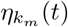 and 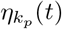 qualitatively follow the intrinsic noise profile. Indeed, the extrinsic fluctuations uniformly affect gene expression during the activation dynamics by shifting the total noise in protein level to higher values.

Note that the predicted steady-state levels of fluctuations are all very similar in this setting. In principle, fluctuations on different production rates can produce slightly different levels of steady-state protein noise. In fact, fluctuations of the translation rate (and thus of the protein burst size) lead to higher protein noise, as the comparison between their analytical expressions in Eq. 11 and Eq. S18 shows. However, the additional noise term has a factor 1/*p*_*ss*_ that makes it negligible for highly expressed genes. Analogously, as explained in details in the SI File section S1.3, the estimated value of 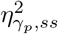 for fluctuations of the degradation rate has a particularly compact form when the timescale of extrinsic fluctuations approximately matches the intrinsic one (as in the example considered):

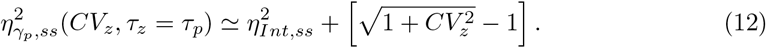

The Taylor expansion of the extrinsic contribution for small 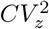 is simply 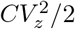, precisely as in the case of fluctuations on production rates (Eq. (11)).

On the other hand, the protein noise dynamics *η*(*t*) displays a qualitatively different and non-monotonic trend for fluctuations of degradation rates (*γ*_*m*_ or *γ*_*p*_). In particular, extrinsic fluctuations on *γ*_*p*_ do not affect the noise dynamics at short times. The total noise is simply dominated by the intrinsic part as the simulations (blue circles) precisely lay on the intrinsic noise theoretical prediction (grey line) for *t < τ*_*p*_ in Fig. 2C. As long as the number of proteins is small, the degradation term is negligible and proteins accumulate approximately linearly with a slope that does not depend on *γ*_*p*_. This can be easily proven by considering the Taylor expansion for *t* → 0 of ⟨*p*(*t*)⟩ when *γ*_*m*_ ≫ *γ*_*p*_:

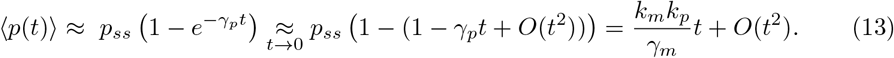

The contribution of *γ*_*p*_ on the protein dynamics, and thus also on the fluctuations, becomes significant only at sufficiently long times. In a similar way, the impact of the cellular factor on *γ*_*m*_ is negligible at the beginning of the simulation and grows in time with ⟨*m*(*t*)⟩.

The generalization of this simple argument allows us to roughly quantify the impact of extrinsic fluctuations at any time through the factorization explained in the Methods section. When extrinsic fluctuations affect the protein dilution rate *γ*_*p*_, our approximate prediction for the time-evolution of protein noise is

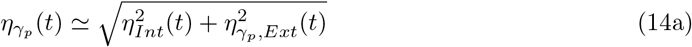

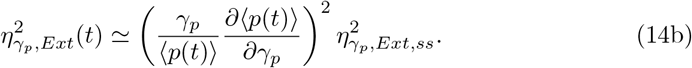

This approximation is displayed as a dashed cyan line in Fig. 2C and well captures the results of exact simulations.

In the case of fluctuations of *γ*_*m*_ we simply tested the validity of the approach summarized in Eq. (8) without explicitly calculating the noise at steady state. Thus, the experimental value at steady state was used to obtained the dynamics represented in Fig. 2C as a dashed magenta line. Also in this case the approximation captures the empirical trend.

The minimum values of 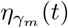 and 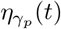 mark the times at which extrinsic fluctuations start to significantly affect the expression noise. As intuitively expected, this happens approximately at the corresponding molecule lifetimes since they set the timescales of the approach of the steady state. In particular, the transition is at a time ≃ 1 (in units of *τ*_*p*_) in Fig. 2C for noise on *γ*_*p*_, and earlier for noise on *γ*_*m*_ since the mRNAs approaches the steady state quickly. Fig. 2 refers to a particular choice of *CV*_*z*_ and *τ*_*z*_=*τ*_*p*_ (matched timescales), but the trends are robust with respect to the values of *CV*_*z*_ and *τ*_*z*_, as reported for example in Fig.S2.

A relevant implication of this analysis is that the simple observation of the dynamics of protein noise, which is experimentally accessible, can clearly distinguish between alternative sources of gene-expression variability, even if the parameter values are not known.

### Linear increase of protein noise with the strength of extrinsic fluctuations

This section explores in detail the role of the strength of extrinsic fluctuations in shaping the single-cell expression dynamics. Fig. 3A reports the protein noise *η*(*t*) as a function of the extrinsic noise *CV*_*z*_. During the early stages of the protein activation dynamics (time *t* = 0.4*τ*_*p*_) the intrinsic component of expression noise (grey line) is significant, and in particular it is the only contribution to the total noise when extrinsic fluctuations act on *γ*_*p*_. For sufficiently long times, the intrinsic noise becomes less relevant, and the expression variability starts to increase linearly with the strength of extrinsic fluctuations. The proportionality coefficient depends on the fluctuating parameter and on the time of the measurement. The figure refers to the biologically relevant situation of approximately matched timescale *τ*_*z*_=*τ*_*p*_. In this case, the cellular factor *z* can represent a protein whose fluctuations affect the production or the degradation of a protein of interest *p* with a similar average lifetimes (for example set by the cell doubling time). In these conditions, *η* reaches a single steady-state level for long times (*t*=20*τ*_*p*_) for any choice of the fluctuating parameter, as displayed by the overlap of the different curves in the right panel of Fig. 3A.

**Fig 3.**
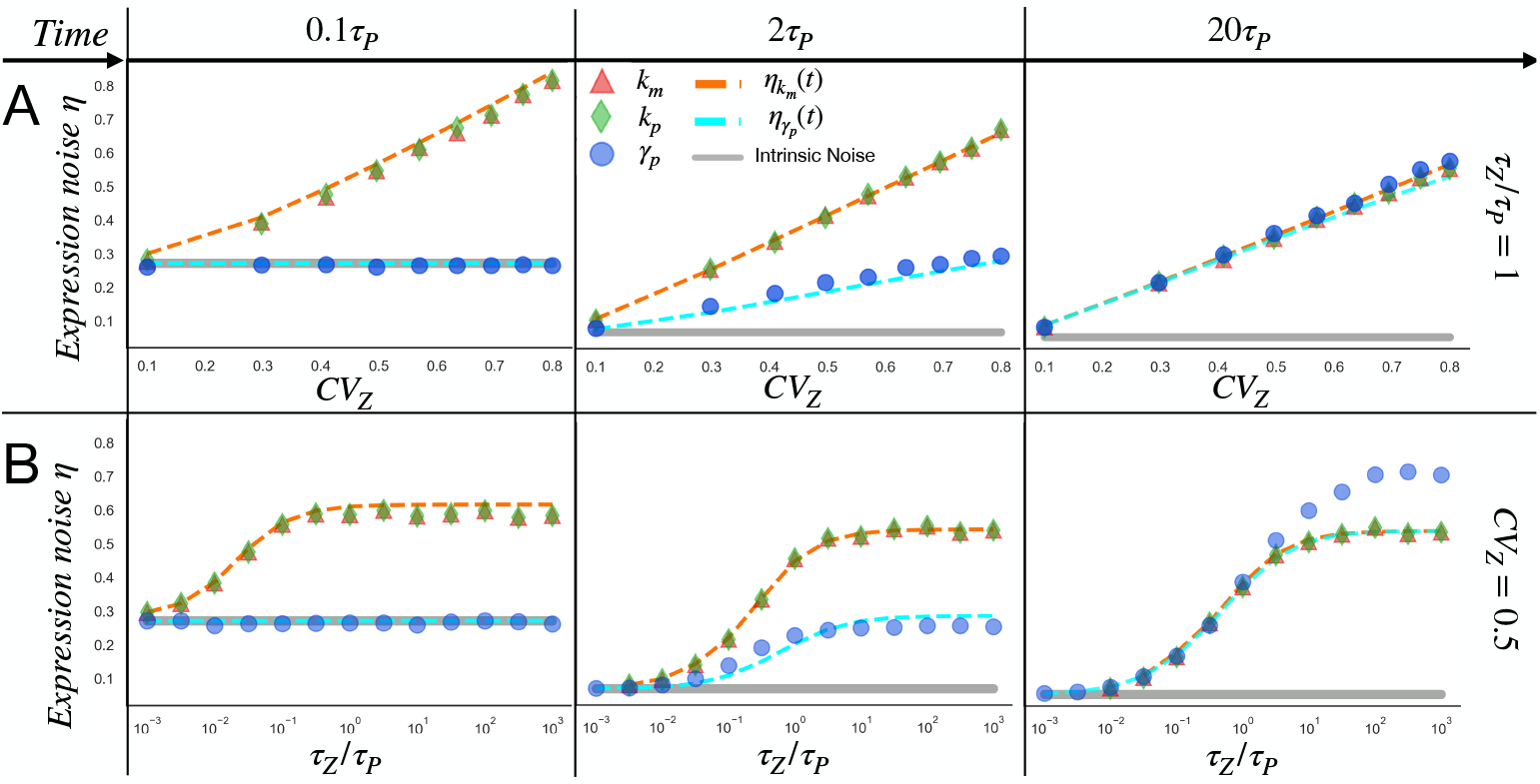
Functional dependence of protein variability on the extrinsic noise strength and timescale. The protein noise is reported as a function of the strength (A) and timescale (B) of extrinsic fluctuations. Different curves correspond to fluctuations of different expression parameter (*k*_*m*_*, k*_*p*_ or *γ*_*p*_) as explained in the legend. The analysis is reported at three different times in the activation dynamics, representing the early stage, an intermediate stage and essentially the steady state. Dashed lines represent the theoretical predictions of Eq. (10) (orange) and Eq. (14) (cyan) that well capture the results of Gillespie simulations. To investigate the role of *CV*_*z*_ (A), the timescale of *z*(*t*) is fixed. In this example it matches the timescale of *p*(*t*), i.e., *τ*_*z*_ = *τ*_*p*_. On the other hand, to explore the role of the timescale *τ*_*z*_ (B), the coefficient of variation of the cellular factor has been held constant to *CV*_*z*_ = 0.5.

The dashed lines in Fig. 3A correspond to our theoretical predictions for 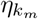 and 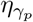, i.e., respectively Eq. (10) and Eq. (14), and the plots show the agreement with simulation results. Again, we are comparing systems that converge to very similar steady state levels of noise. The difference grows in principle with *CV*_*z*_, but it is negligible within our range of parameter exploration. This can be shown by comparing the Taylor expansion of Eq. 12 for small 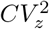 to the extrinsic contribution in Eq. 11.

### Sigmoidal dependence of protein noise on the timescale of extrinsic fluctuations

Fig. 3B displays the role of the timescale of extrinsic fluctuations in determining the time evolution of protein variability. The consequences of adding stochasticity through the coupling with an extrinsic factor *z*(*t*) are minimal when its autocorrelation time is much smaller than the duration of a typical intrinsic fluctuation of *p*(*t*), which is set by the protein lifetime. In fact, when these two timescales are well separated, i.e., *τ*_*z*_ ≪ *τ*_*p*_ or equivalently *γ*_*z*_ ≫ *γ*_*p*_, extrinsic fluctuations are simply averaged out in the protein dynamics and the extrinsic component is negligible. Indeed, all noise curves start from the intrinsic noise baseline (gray line) for relatively fast *z* fluctuations in Fig. 3B.

In the opposite setting (i.e., *τ*_*z*_ ≫ *τ*_*p*_ or *γ*_*z*_ ≪ *γ*_*p*_), the extrinsic factor evolves very slowly in time. Therefore, every cell in the population has a specific random value for the fluctuating parameter that is essentially frozen during the protein dynamics. The system is essentially subjected to what is typically called *quenched disorder* in statistical physics. Around *τ*_*z*_/*τ*_*p*_ *>* 10 the system enters into this “quenched regime” and the noise *η* saturates at a value that only depends on the level *CV*_*z*_ of the quenched noise. The presence of these two regimes makes the dependence on the extrinsic-noise timescale sigmoidal and the crossover between the two regimes is at *τ*_*z*_ ≃ *τ*_*p*_ as intuitively expected.

As mentioned before, at the beginning of the dynamics (*t*=0.04*τ*_*p*_), intrinsic noise is dominant when the fluctuating parameter is *γ*_*p*_ (blue circles), and thus the extrinsic contribution to *η* is negligible independently of its timescale. This is not the case for fluctuations in production parameters. The discrepancy is well capture by our analytical predictions (dashed lines). For longer times, such as *t*=2*τ*_*p*_, the cell-to-cell variability continuously moves from a dominant intrinsic noise for short living extrinsic fluctuations (*τ*_*z*_ ≪ *τ*_*p*_) to the opposite situation in the quenched regime (*τ*_*z*_ ≫ *τ*_*p*_), since in this example *CV*_*z*_ ≫ *η*_*Int,ss*_.

At the steady state (*t*=20*τ*_*p*_ in the figure) and in the quenched regime, the gene expression variability is essentially set by the distribution of the extrinsic factor *z*. Our analytical predictions converge for any fluctuating parameter to a single steady-state value given by the expression

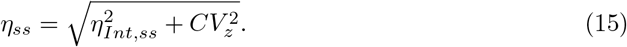

However, while the analytical expression seem generally accurate, the noise exceeds the prediction in simulations with *γ*_*p*_ as a fluctuating parameter, indicating the presence of a noise amplification which is not captured by our analytical approximations.

### Fluctuations of protein dilution rate after a transcriptional block enhances expression noise

The analysis has focused so far on the gene activation dynamics. This section studies the time-evolution of protein noise in the case of a sudden transcriptional repression. We consider a gene expressed at steady-state level whose transcription rate goes to zero at time zero, and we analyze the following protein dynamics when extrinsic noise affects different parameters. Again a mathematically controlled comparison (defined in the Methods section) will be used: the system always start from the same average steady state level as in Fig. 4A. Fluctuations acting on different parameters do not alter the average protein decay dynamics (Fig. 4A), which is well described by Eq. (3). Analogously, the initial level of fluctuations is given by Eqs. (12) and (11) and is approximately independent on which is the fluctuating parameter in this setting, as it is also shown by the overlap of the protein distributions at the initial time in Fig. 4B. Nevertheless, clear differences in the protein probability densities appear as the switch-off dynamics unfolds (Fig. 4B). In particular, extrinsic fluctuations predominantly on the protein degradation rate leads to high-variance protein distributions.

**Fig 4.**
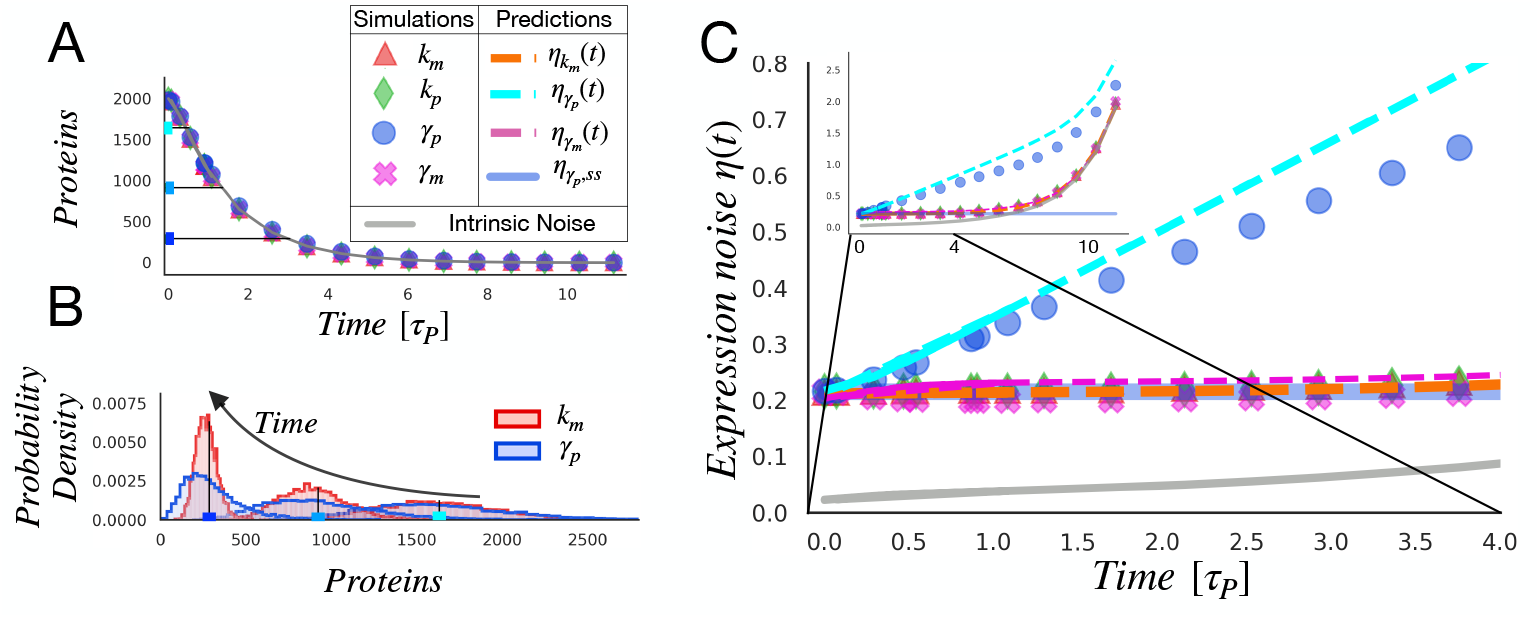
Time-dependent cell-to-cell variability after a sudden transcriptional block. In analogy with Fig. 2, throughout the four alternative settings, we maintain the inactivation dynamics and we fix the properties of the extrinsic noise. A) inactivation dynamics is almost not influenced by extrinsic noise independently of its source, making the controlled comparison method redundant for this particular choice of *CV*_*z*_=0.3, *τ*_*z*_/*τ*_*p*_=1, ⟨*z*⟩=1000 cellular factors. B) The main differences between proteins probability densities in case of fluctuations of *k*_*m*_ or *γ*_*p*_ are appreciable during the intermediate and the final stage of the transient. C) The time-evolution of expression variability in case of fluctuations of production rates or mRNA’s degradation rate is very similar to the one that we observe in the absence of the source of extrinsic noise (grey continuous line). When fluctuations act on *γ*_*p*_ their effect is to enhance expression variability, although the general monotonicity is maintained.

As the expression level progressively approaches zero, the intrinsic contribution grows exponentially, following the grey continuous line in the inset of Fig. 4C. After the transcriptional block, the number of mRNAs decays with a timescale set by *τ*_*m*_, which is typically short with respect to the protein lifetime. Therefore, ⟨*m*(*t*)⟩ quickly goes to zero and protein degradation is the only possible reaction left. This explains why the expression noise profiles are equivalent to the intrinsic noise prediction with the exception of 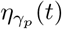, i.e., when extrinsic fluctuations predominantly affects the protein lifetime.

Our theoretical predictions for *η*(*t*) (dashed lines of Fig. 4C) derive from the simple argument of Eq. 8, which when applied in this context gives:

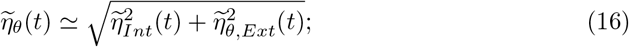

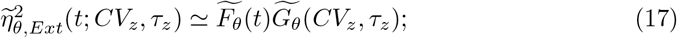

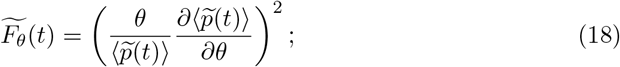

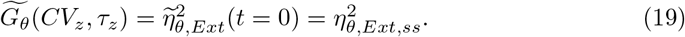

To determine the role of 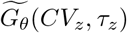, the limit lim_*t*⟶0_ 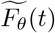 has to be considered, which correspond to the initial steady state. In the case of fluctuations of *k*_*m*_*, k*_*p*_ or *γ*_*m*_, the term 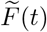 does not depend on time. Therefore, the noise dynamics is uniformly affected by extrinsic noise. *η*(*t*) remains approximately constant until the intrinsic contribution becomes dominant (around *t* ≃ 7*τ*_*p*_) and makes the noise grow exponentially.

The trend of 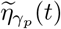, i.e., for a fluctuating protein degradation rate, is qualitatively different and well predicted by our analytical estimations. As long as the intrinsic contribution is negligible, protein noise grows linearly with time as 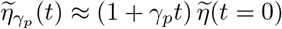. For sufficient long times, *p*(*t*) approaches zero and again the noise diverges because of the intrinsic contribution.

The fluctuation trends in the gene deactivation dynamics do not have qualitatively different behaviors as in the case of gene activation, there is a general increase in protein noise [55]. However, an extrinsic noise that mainly affects the degradation rate significantly increases the protein fluctuations immediately after the transcriptional block. On the other hand, it is necessary to wait way longer then the protein lifetime to see a considerable fold in protein noise if the production rates are fluctuating. An estimate of the protein lifetime can be easily extracted from the average protein dynamics after the transcriptional block. In fact, the average protein value follows Eq. 3, which is simply an exponential decay with exponent given by the protein lifetime if the protein is stable with respect to the mRNA. These observations can be used in time-coarse experiments to identify the main form of extrinsic noise in the system in analysis.

### The interplay between multiple fluctuating parameters

One single parameter could be predominantly affected by extrinsic fluctuations as we have assumed so far. For example, if fluctuations in ribosome concentration are the main source of extrinsic noise, as it has been hypothesized in fast-growing bacteria [41], the translation rate would be the main fluctuating parameter in a corresponding model of stochastic gene expression. However, more general variability in cell physiology can affect multiple expression parameters in different ways. For example, growth rate is coupled to ribosome concentration as well as to cell volume, and thus its fluctuations can influence translation rate as well as protein dilution. Therefore, this section explores the consequences of extrinsic fluctuations affecting two different expression parameters with comparable intensity. In particular, we focus on the illustrative example of fluctuations in the transcription rate and in the protein dilution/degradation rate, and we describe their possible interactions in defining the final level of protein noise.

Previous studies have shown that extrinsic fluctuations can combine constructively or destructively at the steady state depending on their action on parameters [39, 40]. Here, we extend the analysis to the out-of-equilibrium dynamics.

More specifically, a single extrinsic factor *z* can simultaneously set the fluctuations of the two different parameters in a positively or negatively correlated fashion. Alternatively, two independent sources of noise (*z*_1_ and *z*_2_) can individually affect the two different parameters making their fluctuations uncorrelated. While these represent the more clear-cut scenarios, a specific biological system could present multiple sources of extrinsic noise and thus intermediate and more nuanced situations.

When two sources of noise independently affect the transcription and degradation rates, a simple additive combination can be observed. In other words, the total extrinsic noise is well described by the sum of the extrinsic contributions we previously calculated with a single fluctuating parameter. Therefore, the total protein noise can be simple written as

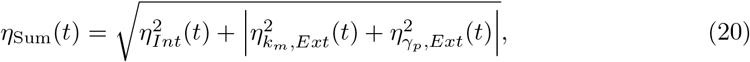

where 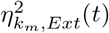 and 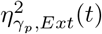 are the expressions calculated for *k*_*m*_ or *γ*_*p*_ as the only fluctuating parameter. Fig. 5 shows that this analytical prediction fits rather well the simulation results (green dots).

**Fig 5.**
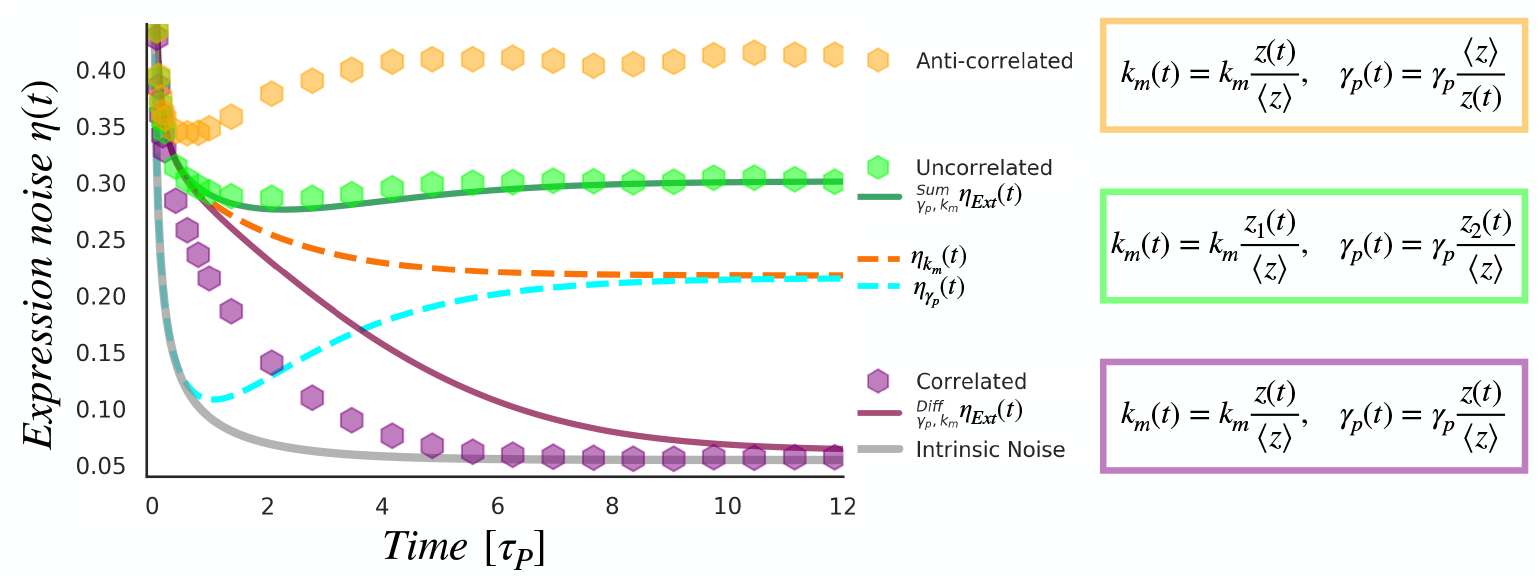
Simultaneous fluctuations of k_m_ and *γ*_p_ can combine constructively or destructively during transient dynamics. We consider the combined action of extrinsic alterations on the transcription rate and the dilution rate. The fluctuations are either uncorrelated and generated by individual sources of stochasticity, *z*_1_(*t*) and *z*_2_(*t*), (in green), or generated by the same source, *z*(*t*), affecting the parameters in a correlated (purple) or anti-correlated way (yellow). The simulation specifications are: ⟨*z*_1_⟩= ⟨*z*_2_⟩= ⟨*z*⟩=1000 cellular factors, 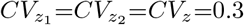 and 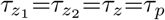.

When a single extrinsic factor *z* sets the two parameters in a correlated way, a positive fluctuation in *z* increases the protein production rate but, at the same time, also boosts the degradation rate. Therefore, the two fluctuations combine destructively, and the resulting extrinsic contribution is approximately the difference between the two noise contributions. The total protein noise can thus be described by

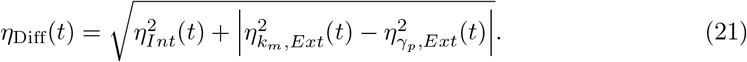

This expression is plotted as a continuous purple line in Fig. 5 and captures the behavior of the corresponding simulations (purple dots).

On the contrary, when a single source of stochasticity affects *k*
_*m*_ and *γ*_*p*_ in an anti-correlated way, their corresponding noise contributions combine constructively, so that the total noise exceeds *η*_Sum_(*t*).

The non-monotonous trend of protein noise, which is the hallmark of dominant extrinsic fluctuations in *γ*_*p*_ as shown in Fig 2, is thus conserved also if another parameter fluctuates with an equivalent variance, as long as the two contributions do not combine destructively.

## Discussion

From the first quantitative single-cell experiments with clonal bacterial populations, the substantial cell-to-cell variability in gene expression was evident, and could be traced back to two distinct origins, i.e., intrinsic and extrinsic noise [36, 38]. Experimentally, the dual-color experiment allows to disentangle the two noise factors [38], although their interplay can actually be complex and not trivial to interpret [19]. Large-scale studies showed the major relevance of the extrinsic contribution, especially for sufficiently expressed genes [35, 41, 42]. In parallel, theoretical analysis have been proposed on the consequences of extrinsic noise on protein distributions at steady state [39, 40]. However, understanding the dominant biological sources of extrinsic noise in a specific organism, and thus the expression rates mostly affected by these general cellular factors, has proven to be a difficult task. A major problem is that factors related to cell physiology such as metabolism or cell cycle, which are known to have substantial cell-to-cell variability [20, 56, 57], can affect multiple steps in gene expression from the gene copy number to the translation rate. This makes hard to hypothesize what rates are actually predominantly fluctuating and thus to build appropriate mathematical models.

The observation that equilibrium distributions could not be sufficient to fully characterize the noise sources was previously recognized in an analysis on the role of RNA degradation fluctuations [58]. Analogously, the idea that the molecule dynamics can provide additional information on the dominant source of noise has been exploited to distinguish between alternative intrinsic noise models [55, 59]. This paper extends these intuitions and provides a detailed theoretical analysis of the possible scenarios in which different parameters are predominantly coupled with an extrinsic noise source. More specifically, we characterized the dynamics of the average protein level and of its fluctuations out-of-equilibrium, showing that the dynamics specifically presents signatures of the extrinsic noise source. Current experimental techniques based on fluorescence time-lapse microscopy [60, 61], potentially coupled with microfluidic devices to keep cells in a controlled environment for many generations [61, 62], give access to the expression dynamics at the single-cell level when a reporter gene is induced or suppressed. Therefore, our results can be directly compared with experimental trends, providing a simple tool to better understand the dominant sources of extrinsic noise by pinpointing the fluctuating expression parameters. We also explored in detail the parameter space, providing analytical estimates of how the picture can quantitatively change as a function of the timescale and strength of extrinsic fluctuations.

We used a simple and general modelling framework to keep it amenable of analytical calculations and applicable to different biological systems. The actual cellular mechanisms giving rise to extrinsic noise are only phenomenologically captured by a generic “cellular factor” *z* described by a three-parameter distribution of which we varied the parameter values. An alternative (and complementary) approach would be to start from specific mechanistic descriptions of global cellular factors, for biological systems in which are available, and explore their impact on noise in gene expression. For example, in fast growing bacteria quantitative descriptions of the cell cycle [63, 64] and of the “laws” governing the global resource partitioning in the cell [43, 65, 66] have been proposed. These basic aspects of cell physiology greatly contribute to gene expression rates and their fluctuations by setting the cell volume, protein dilution, as well as the concentration of key enzymes [67]. Along this line, recent work focused on the effect of cell cycle and growth rate variability on gene expression noise [30, 32, 68]. It would be interesting to analyze the relation between specific mechanistic models of the extrinsic noise source and the statistics of the general factor *z* in our approach, in order to apply our analytical results and predictions for the out-of-equilibrium dynamics to more detailed biological descriptions.

While this work focuses on the dynamics of an isolated gene, a natural extension will be to consider more complex regulatory interactions, such as the ubiquitous circuits of auto-regulation, feed-forward loops and regulatory cascades. Noise propagation through transcriptional regulation is a known substantial source of extrinsic noise [33, 34], and thus the dynamical fluctuation properties here described could in principle change in presence of regulatory circuits.

Finally, the presence of extrinsic noise can have relevant consequences on the timing precision of genetic circuits. Models of stochastic timing in gene expression have typically focused on the consequences of intrinsic noise [69–71]. However, characterizing the out-of-equilibrium dynamics with extrinsic noise, the results presented here can be easily used to derive estimates of first-passage-time distributions in presence of different types of extrinsic fluctuations.

## Supporting information

### SI File

The supplementary information file contains: the detailed description of our method to implement a source of extrinsic noise in the standard stochastic model of gene expression; the derivation of the analytical predictions presented throughout the Results section; the specification of the parameters used during the simulations.

## Author contributions

Conceptualization: MO; methodology: AS, MO and MB; formal analysis and software: MB; original draft preparation: MB and MO; review and editing: MB, AS, MC and MB. All authors have read and agreed to the published version of the manuscript.

## Acknowledgments

We would like to thank all the members of the INFN BioPhys group of the University of Turin for the useful discussions.

