## Supplementary material for "Out-of-equilibrium gene expression fluctuations in presence of extrinsic noise": SI File

#### S1 Modelling extrinsic noise

The basic model of gene expression consists of four possible events that occur randomly at exponentially-distributed time intervals, with rates that are constant in absence of an extrinsic source of noise. The discrete changes in the mRNA and protein populations due to the four events are listed in Tab. S1. The third column shows the event propensity function that determines how often an event occurs.

| Event | Population reset | Propensity function ($f$) |
|---|---|---|
| mRNA birth | $m(t) \to m(t) + 1$ | $k_m$ |
| mRNA death | $m(t) \to m(t) - 1$ | $m(t)\gamma_m$ |
| Protein birth | $p(t) \to p(t) + 1$ | $m(t)k_p$ |
| Protein death | $p(t) \to p(t) - 1$ | $p(t)\gamma_p$ |

Supplementary Table S1: The standard stochastic model of gene expression.

In the often-valid limit of short-living mRNAs with respect to proteins ($\gamma_m \gg \gamma_p$), the model can be approximated by a simple *bursty expression* model: bursts of protein production arrive at a constant rate $k_m$ (as in the Poisson transcription process) and the burst size is given by the number of proteins produced by a single mRNA $b_p$. $b_p$ is a random variable following a geometric probability distribution with mean burst size $B_p = k_p/\gamma_m$ [1].

For the above model, the time derivative of the expected value of any differentiable function $\varphi(m,p)$ is given by

$$\frac{d\langle\varphi(m,p)\rangle}{dt} = \langle \sum_{Events} \Delta\varphi(m,p) \times f(m,p)\rangle \tag{S1}$$

where $\Delta\varphi(z,m,p)$ is the change in $\varphi(z,m,p)$ when an event occurs and $f(m,p)$ is the event propensity function [2]. In particular, the moments of $p(t)$ can be directly obtained from the corresponding Chemical Master Equation [3, 4]. For each positive integer $n$, the time evolution of the expected value of $p(t)^n$ is given by

$$\frac{d\langle p(t)^n\rangle}{dt} = \langle G(p)\rangle, \quad n \in \{0,1,2,...\}, \tag{S2}$$

$$G(p) := \sum_{j=0}^{\infty} k_m \mathbb{P}(b_p = j)[(p+j)^n - p^n] + \gamma_p p[(p-1)^n - p^n]. \tag{S3}$$

$\mathbb{P}(b_p = j)$ is the probability of having a burst of $j$ protein molecules. From this expression, we can write the equations for the dynamics of the first two moments as

$$\frac{d\langle p\rangle}{dt} = k_m B_p - \gamma_p\langle p\rangle \tag{S4a}$$

$$\frac{d\langle p^2\rangle}{dt} = \gamma_p(\langle p\rangle - 2\langle p^2\rangle) + k_m\langle b_p^2\rangle + 2k_m\langle p\rangle B_p, \tag{S4b}$$

where we substitute the moments of the burst size geometrical distribution, i.e., $\langle b_p^2\rangle = 2\langle b_p\rangle^2 + \langle b_p\rangle = 2B_p^2 + B_p$.

By considering the steady state of Eqs. (S4), the protein mean and noise levels can be calculated as

$$p_{ss} = \frac{k_m}{\gamma_p} B_p, \quad \eta^2 = \frac{1}{p_{ss}}(1 + B_p) := \eta^2_{Int,ss} \tag{S5}$$

We now introduce extrinsic noise in the model using an extrinsic factor with copy number $z(t)$ whose stochastic dynamics is modeled as a bursty process with constant production rate $k_z$ and degradation rate $\gamma_z$. A production event generates a geometrically distributed burst of $b_z$ molecules with average burst size $B_z$ following

$$\mathbb{P}\{b_z = i\} = \left(1 - \frac{1}{B_z}\right)^{i-1} \frac{1}{B_z}, \quad i = 1,2,3,... \tag{S6}$$

The advantage of this phenomenological description is that the extant and the timescale of fluctuations in $z(t)$ can be independently modulated by tuning $k_z$, $\gamma_z$ and $B_z$.

We always consider the cellular factor stochastic process $z(t)$ to be at the steady state characterized by a mean value $z_{ss}$ and $CV_z^2$. For such a process, the steady-state mean $z_{ss}$, the coefficient of variation squared $CV_z^2$ and the autocorrelation function $R_z(\delta t)$ are given by the following relationships:

$$z_{ss} = \frac{k_z}{\gamma_z} B_z \tag{S7}$$

$$CV_z^2 = \frac{1}{z_{ss}}(1 + B_z) \tag{S8}$$

$$R_z(\delta t) = e^{-\gamma_z \delta t} \tag{S9}$$

The timescale associated with degradation $\gamma_z$ sets the steady-state autocorrelation time of the bursty birth-death process $z(t)$. $1/\gamma_z$ describes the average lifetime of a typical fluctuation, as well as the average time separating such fluctuations. Therefore, $CV_z$ and $\tau_z$ respectively represent the *extent* and the *timescale* of fluctuations induced in certain parameters of the model.

For any parameter $\theta$ of the system, extrinsic fluctuations were implemented by modifying the propensity of the respective reaction such that at any moment the time-dependent rate was $\theta(t) = \theta \frac{z(t)}{\langle z \rangle}$, with $\langle \theta(t) \rangle = \theta$ and $CV_\theta^2 = CV_z^2$.

#### S1.1 Transcription burst frequency fluctuations

Fluctuations in the extrinsic factor level may impact protein synthesis via its transcription rate, as formalized in Tab. S2. This leads to a system of coupled bursty birth-death processes.

The time-derivative of the expected value of any differentiable function $\varphi(z, m, p)$ is given by:

$$\frac{d\langle \varphi(z, m, p) \rangle}{dt} = \langle \sum_{Events} \Delta\varphi(z, m, p) \times f(z, m, p) \rangle \tag{S10}$$

| Event | Population reset | Propensity function ($f$) |
|---|---|---|
| Cellular factor birth | $z(t) \to z(t) + i$ | $k_z \mathbb{P}\{b_z = i\}$ |
| Cellular factor death | $z(t) \to z(t) - 1$ | $z(t)\gamma_z$ |
| mRNA birth | $m(t) \to m(t) + 1$ | $\frac{z(t)}{z_{ss}} k_m$ |
| mRNA death | $m(t) \to m(t) - 1$ | $m(t)\gamma_m$ |
| Protein birth | $p(t) \to p(t) + 1$ | $m(t)k_p$ |
| Protein death | $p(t) \to p(t) - 1$ | $p(t)\gamma_p$ |

Supplementary Table S2: An effective model of gene expression in the presence of a source of extrinsic noise.

The statistical moments of this joint process evolve as:

$$\frac{d\langle p(t)^{n_1} z(t)^{n_2}\rangle}{dt} = \langle G(p, z)\rangle, \quad n_1, n_2 \in \{0, 1, 2, ...\} \tag{S11}$$

$$\begin{aligned} G(p, z) := &\sum_{j=0}^{\infty} \frac{z(t)k_m}{z_{ss}} \mathbb{P}(b_p = j)[(p + j)^{n_1} z^{n_2} - p^{n_1} z^{n_2}] + \\ &\sum_{i=1}^{\infty} k_z \mathbb{P}(b_z = i)[p^{n_1}(z + i)^{n_2} - p^{n_1} z^{n_2}] + \\ &\gamma_p p[(p - 1)^{n_1} z^{n_2} - p^{n_1} z^{n_2}] + \gamma_z z[p^{n_1}(z - 1)^{n_2} - p^{n_1} z^{n_2}] \end{aligned} \tag{S12}$$

[3, 4]. Substituting the appropriate values of $n_1$ and $n_2$ yields the time evolution of the first and second-order moments of $p(t)$ and $z(t)$

$$\frac{d\langle z\rangle}{dt} = k_z B_z - \gamma_z \langle z\rangle \tag{S13a}$$

$$\frac{d\langle p\rangle}{dt} = \frac{\langle z\rangle k_m B_p}{z_{ss}} - \gamma_p \langle p\rangle \tag{S13b}$$

$$\frac{d\langle z^2\rangle}{dt} = k_z \langle b_z^2\rangle + 2k_z \langle z\rangle B_z + \gamma_z(\langle z\rangle - 2\langle z^2\rangle) \tag{S13c}$$

$$\frac{d\langle p^2\rangle}{dt} = \frac{\langle z\rangle k_m \langle b_p^2\rangle}{z_{ss}} + \frac{2k_m \langle pz\rangle B_p}{z_{ss}} + \gamma_p(\langle p\rangle - 2\langle p^2\rangle) \tag{S13d}$$

$$\frac{d\langle pz\rangle}{dt} = k_z B_z \langle p\rangle + \frac{k_m B_p \langle z^2\rangle}{z_{ss}} - \gamma_p \langle pz\rangle - \gamma_z \langle pz\rangle \tag{S13e}$$

Solving the above system of differential equations assuming that $p(0) = 0$ and that the extrinsic factor is at steady-state at time $t = 0$ provides both the average $p(t)$ and protein noise level over time:

$$p_{k_m}(t) = \frac{\langle z \rangle}{z_{ss}} \langle p(t) \rangle \Rightarrow p_{k_m}(t_{ss}) = \frac{\langle z \rangle}{z_{ss}} \frac{k_m B_p}{\gamma_p} = p_{ss} \tag{S14a}$$

$$\eta^2_{k_m}(t; CV_z, \tau_z) = \frac{2e^{\gamma_p t}}{(e^{\gamma_p t} - 1)^2} \Bigg\{ \eta^2_{Int,ss} \sinh(\gamma_p t) + \tag{S14b}$$
$$\frac{\gamma_p^2[\cosh(\gamma_p t) - \cosh(\gamma_z t) + \sinh(\gamma_z t)] - \gamma_p \gamma_z \sinh(\gamma_p t)}{\gamma_p^2 - \gamma_z^2} CV_z^2 \Bigg\}$$

Since $\langle z \rangle = z_{ss}$, the average protein dynamics is not affected by fluctuations of the gene transcription rate.

The steady-state noise levels are now given by:

$$\eta^2_{k_m,ss} = \frac{1}{p_{ss}}(1 + B_p) + \frac{\gamma_p}{\gamma_p + \gamma_z} CV_z^2 = \eta^2_{Int,ss} + \frac{\gamma_p}{\gamma_p + \gamma_z} CV_z^2 \tag{S15}$$

The first component is the intrinsic one while the second component is due to the extrinsic factor contribution [5]. We stress that individual $k_m$ or $k_p$ fluctuations do not impact the steady-state mean protein levels, given by Eq. (S5).

#### S1.2 Translation rate fluctuations

To model noise in the protein translation rate, we modify $k_p$ to $k_p z(t)/z_{ss}$. This will cause fluctuations of the protein burst size:

$$\langle b_p \rangle = \sum_{j=0}^{\infty} \mathbb{P}(b_p = j) j = \frac{z(t)}{z_{ss}} B_p \tag{S16a}$$

$$\langle b_p^2 \rangle = \sum_{j=0}^{\infty} \mathbb{P}(b_p = j) j^2 = 2 \left( \frac{z(t)}{z_{ss}} B_p \right)^2 + \frac{z(t)}{z_{ss}} B_p \tag{S16b}$$

The time evolution of moments is then given by:

$$\frac{d\langle z\rangle}{dt} = k_z B_z - \gamma_z \langle z\rangle \tag{S17a}$$

$$\frac{d\langle p\rangle}{dt} = \frac{k_m \langle z\rangle B_p}{z_{ss}} - \gamma_p \langle p\rangle \tag{S17b}$$

$$\frac{d\langle z^2\rangle}{dt} = k_z \langle b_z^2\rangle + 2k_z \langle z\rangle B_z + \gamma_z(\langle z\rangle - 2\langle z^2\rangle) \tag{S17c}$$

$$\frac{d\langle p^2\rangle}{dt} = 2k_m \frac{B_p \langle pz\rangle}{z_{ss}} + 2k_m \frac{B_p^2 \langle z^2\rangle}{z_{ss}^2} + k_m B_p - 2\gamma_p \langle p^2\rangle + \gamma_p \langle p\rangle \tag{S17d}$$

$$\frac{d\langle pz\rangle}{dt} = k_z B_z \langle p\rangle + k_m \frac{B_p \langle z^2\rangle}{z_{ss}} - \gamma_z \langle pz\rangle - \gamma_p \langle pz\rangle \tag{S17e}$$

Once again the average protein dynamics is not affected by the extrinsic fluctuations. The steady-state noise levels are now given by:

$$\begin{aligned} \eta^2_{k_p,ss} &= \frac{1}{p_{ss}}(1 + B_p) + \frac{\gamma_p}{\gamma_p + \gamma_z} CV_z^2 + \frac{B_p}{p_{ss}} CV_z^2 = \\ &= \eta^2_{Int,ss} + \frac{\gamma_p}{\gamma_p + \gamma_z} CV_z^2 + \frac{B_p}{p_{ss}} CV_z^2 = \\ &= \eta^2_{k_m} + \frac{B_p}{p_{ss}} CV_z^2 \end{aligned} \tag{S18}$$

The predicted steady-state protein noise in case of protein burst size fluctuations $\eta^2_{k_p,ss}$ exceeds that caused by transcriptions rate fluctuations of $B_p CV_z^2/p_{ss}$. Since in our range of exploration ($B_p = 5$,$p_{ss} = 2000$ and $CV_z \in [0.1, 0.8]$) the contribution due to $B_p CV_z^2/p_{ss}$ is minimal, we assume $\eta_{k_p} \simeq \eta_{k_m}$ and we did not explicitly reported the trend $\eta_{k_p}(t)$ in the figures of the main text.

#### S1.3 Protein dilution rate fluctuations

To model noise in the decay rate, we modify $\gamma_p$ to $\gamma_p z(t)/z_{ss}$. The time evolution of moments is then given by [3, 4]:

$$\frac{d\langle z\rangle}{dt} = k_z B_z - \gamma_z \langle z\rangle, \tag{S19a}$$

$$\frac{d\langle p\rangle}{dt} = k_m B_p - \gamma_p \frac{\langle pz\rangle}{z_{ss}}, \tag{S19b}$$

$$\frac{d\langle z^2\rangle}{dt} = k_z \langle b_z^2\rangle + 2k_z\langle z\rangle B_z + \gamma_z(\langle z\rangle - 2\langle z^2\rangle) \tag{S19c}$$

$$\frac{d\langle p^2\rangle}{dt} = k_m \langle b_p^2\rangle + 2k_m\langle p\rangle B_p + \frac{\gamma_p(\langle pz\rangle - 2\langle p^2 z\rangle)}{z_{ss}} \tag{S19d}$$

$$\frac{d\langle pz\rangle}{dt} = k_z B_z\langle p\rangle + k_m B_p\langle z\rangle - \frac{\gamma_p\langle pz^2\rangle}{z_{ss}} - \gamma_z\langle pz\rangle \tag{S19e}$$

Note that in this case the moment dynamics is not closed, and the time evolution of lower order moments depends on higher order moments $\langle p^2 z\rangle$ and $\langle pz^2\rangle$. To close moment dynamics we use the derivative-matching closure scheme that is consistent with copy numbers following a lognormal distribution [4]. As per this closure, the higher order moments are approximated as

$$\langle pz^2\rangle \approx \frac{\langle z^2\rangle}{\langle p\rangle}\left(\frac{\langle pz\rangle}{\langle z\rangle}\right)^2 \tag{S20a}$$

$$\langle p^2 z\rangle \approx \frac{\langle p^2\rangle}{\langle z\rangle}\left(\frac{\langle pz\rangle}{\langle p\rangle}\right)^2 \tag{S20b}$$

Substituting the higher order moments in (S19) with their corresponding approximation (S20) results in a closed system of moment dynamics. Solving these equations results in the steady-state protein number and noise levels:

$$p_{\gamma_p}(t_{ss}; CV_z, \tau_z) \simeq \frac{p_{ss}}{2}\left[\sqrt{\left(1+\frac{\tau_z}{\tau_p}\right)^2 + 4\frac{\tau_z}{\tau_p}CV_z^2} + \left(1-\frac{\tau_z}{\tau_p}\right)\right] \tag{S21}$$

$$\eta^2_{\gamma_p,ss}(CV_z,\tau_z) \simeq$$

$$\frac{1}{p_{ss}}(1+B_p) + \left\{\frac{1}{2}\left[\sqrt{\left(1+\frac{\tau_z}{\tau_p}\right)^2 + 4\frac{\tau_z}{\tau_p}CV_z^2} - \left(1+\frac{\tau_z}{\tau_p}\right)\right]\right\} = \tag{S22}$$

$$\eta^2_{Int,ss} + \{\eta^2_{\gamma_p,Ext,ss}\}$$

when the timescale of the fluctuations matches that of the intrinsic fluctuations of the process $p(t)$, the total variability of proteins number at the steady state is the following:

$$\eta^2_{\gamma_p,ss}(CV_z, \tau_z = \tau_p) \simeq \eta^2_{Int,ss} + \left[\sqrt{1 + CV_z^2} - 1\right] \tag{S23}$$

We use this expression of $\eta^2_{\gamma_p,Ext,ss} = \sqrt{1 + CV_z^2} - 1$ to estimate the time-evolution of total noise:

$$\eta_{\gamma_p}(t) \simeq \sqrt{\eta^2_{Int}(t) + \left(\frac{\gamma_p}{p(t)}\frac{\partial\langle p(t)\rangle}{\partial\gamma_p}\right)^2 \eta^2_{\gamma_p,Ext,ss}} \tag{S24}$$

The results of our simulations mostly agree with our analytical prediction Eqs. (S14) and Eqs. (S24), as we can appreciate in Supplementary Fig. S1. However, Eqs. (S24) fails to predict the amplification of the variability added to the system by means of the cellular factor when its typical timescale is longer than that of protein dynamics (in Supplementary Fig. S1B, for long times, the red circles are far above the red continuous line).

#### S1.4 A mathematically controlled comparison.

As reported in the main text, extrinsic fluctuations of the protein dilution rate alter the average protein dynamics. The dependence of the final steady-state level of expression on the properties of the source of the extrinsic noise are given by the approximate expression Eq. (S21). In Supplementary Fig. S2 we can appreciate how our estimate predicts the qualitative trends measured during the simulations. In the light of that premise, it is essential to specify the constraints to put different configurations of the model on equal footing for a fair comparison. Basically, to achieve the same steady-state protein level for any choice of the magnitude and the timescale of extrinsic fluctuations, we slightly change the mean values of $k_m$, $k_p$, $g_m$ to reproduce the appropriate $p_{\gamma_p,ss}(CV_z, \tau_z)$, under the constraint that $\gamma_p$, $B_p$ and $m_{ss}$ are fixed throughout all the different simulation settings.

For any set of values for $CV_z$ and $\tau_z$, the protein expression tends asymptotically to the same equilibrium level independently of the parameter affected by cellular factor fluctuations (Supplementary Fig. S3).

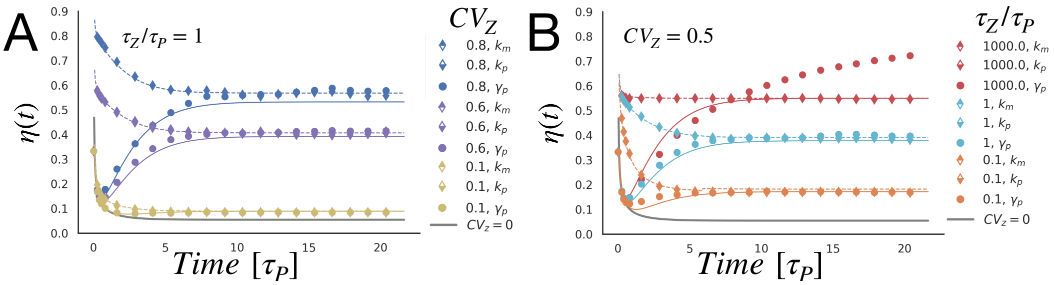


Supplementary Figure S1: **Time evolution of cell-to-cell variability depends on the source of extrinsic noise.** The total gene expression noise is reported as a function of time during the activation dynamics; each color represents a specific value of $CV_z$ (A) or $\tau_z/\tau_p$ (B), while different line-styles correspond to different fluctuating parameters. The continuous grey line represents the total variability that we observe in the system if $CV_z = 0$, i.e. the intrinsic noise. The dashed lines correspond to the theoretical predictions for fluctuations of the transcription rate (Eq. (S14)), they are in good agreement with the results of the simulations. The approximate predictions for expression variability under fluctuations of the protein dilution rate is marked by the continuous lines and correspond to Eq. (S24).

### S2 Stochastic simulation specification

Simulations have been implemented using Gillespie's first reaction algorithm [6]. We simulate the stochastic reactions presented in Fig.1 of the main text and in Tab. S1, with the appropriate changes in the propensities due to the cellular factor. Each data point in the figures is the result of $5 \times 10^3$ trials.

The standard stochastic model of gene expression allows a discrete number of possible configurations, distinguished by the properties of extrinsic noise ($CV_z$, $\tau_z$) and the parameter subjected to it.

We analyse the role of extrinsic fluctuations in shaping expression variability; in particular, we explore the functional dependence of $\eta(t)$ on $CV_z$, and $\tau_z$.

To investigate the role of the extrinsic noise magnitude, we changed the parameter $B_z$ and $k_z$ to span $CV_z$ over a certain range, keeping the value of $\langle z \rangle$ and $\gamma_z$ constant so that the timescale of the process $z(t)$ resulted fixed over all the simulations. Conversely, to investigate the role of the timescale

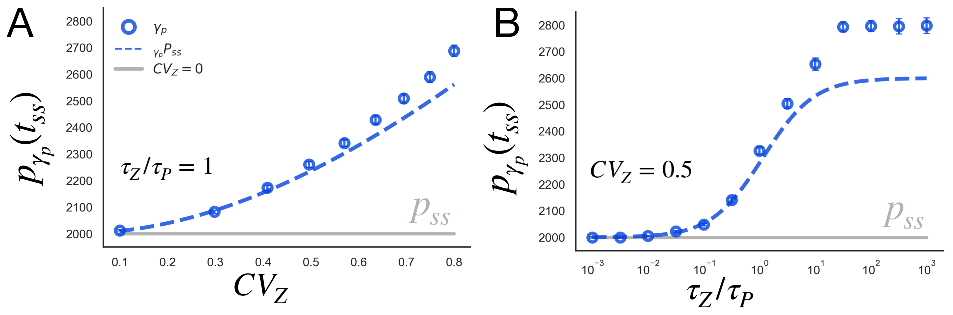


Supplementary Figure S2: **The strength and the timescale of extrinsic fluctuations of the protein dilution rate affect the steady-state level of expression.** Functional dependence of protein steady-state level on the extrinsic noise strength (A) and timescale (B), the dashed lines represent the approximate analytical predictions for $p_{\gamma_p}(t_{ss})$ in case of extrinsic fluctuations acting on the protein dilution rate (Eq. (S21)).

of the colored noise, we properly modulated both $\gamma_z$ and $k_z$ to allow $\tau_z$ to span over a certain range while $\langle z \rangle$ and $CV_z$ were constant. Moreover, the timescale of the extrinsic noise is always expressed in units of $\tau_p$.

We must choose the duration of each simulation. The response times for gene activation and inactivation are both governed by the dilution rate $\tau_p = \ln(2)/\gamma_p$. For proteins that are not actively degraded in the growing cells the response time is equal to one cell generation time [7]. Given this considerations, we set $t_{ss} = 20\tau_p$ as the interruption time for all the simulations since in the absence of extrinsic fluctuations it would largely exceed the time required for the system to reach the steady state.

In the illustrative examples showed in the main text the parameters have the following values:

$$\begin{aligned} k_m &= 8\ mRNA\ min^{-1}; \\ \gamma_m &= 0.2\ min^{-1}; \\ k_p &= 1\ proteins\ (mRNA\ min)^{-1}; \\ \gamma_p &= 0.02\ min^{-1}. \end{aligned} \tag{S25}$$

In this setting, the intrinsic noise is almost negligible because of the low value of the protein burst size $B_p = 5$ proteins mRNA$^{-1}$. Moreover, at the steady-state $m_{ss}$= 40 mRNA and $p_{ss}$=2000 proteins. This values represent

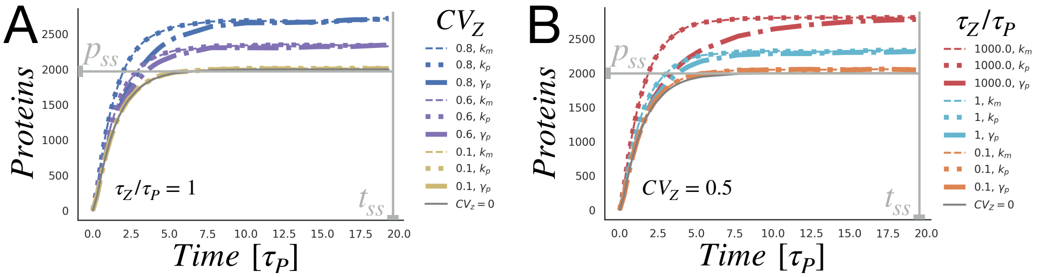


Supplementary Figure S3: **Protein expression tends asymptotically to the same equilibrium level.** At time $t = 0$ transcription begins and the gene is turned on. The average protein level of the simulations approaches the steady-state $p(t_{ss})$, which depends on the characteristics of the extrinsic source of noise. Due to the appropriate tuning of the parameters, when the cellular factor affects $k_m$ or $k_p$ the protein expression tends to the same equilibrium level observed for fluctuations of $\gamma_p$. A) The average protein activation dynamics for $\tau_p/\tau_z = 1$ is reported as a function of time (in units of the response-time), each color represents a specific value of $CV_z$ and different line-styles correspond to different sources of extrinsic noise. B) The average protein activation dynamics for $CV_z = 0.5$ is reported as a function of time, each color represents a specific value of $\tau_z$ (in units of $\tau_p$).

a relatively highly-expressed gene whose expression variability is mainly due to extrinsic fluctuations [8].

[7] Uri Alon. *An Introduction to System Biology.* Chapman & Hall/CRC, Boca Raton FL, 2007.

[8] Yuichi Taniguchi, Paul J. Choi, Gene-Wei Li, Huiyi Chen, Mohan Babu, Jeremy Hearn, Andrew Emili, and X Sunney Xie. Quantifying e. coli proteome and transcriptome with single-molecule sensitivity in single cells. *Science*, 329(5991):533–538, Jul 2010.
